# Structural interplay between DNA-shape protein recognition and supercoiling: the case of IHF

**DOI:** 10.1101/2022.03.31.486518

**Authors:** George D. Watson, Elliot W. Chan, Mark C. Leake, Agnes Noy

## Abstract

The integration host factor (IHF) is a prominent example of indirect readout as it imposes one of the strongest bends on relaxed linear DNA. However, the relation between IHF and torsionally constrained DNA, as occurs physiologically, remains unclear. By using atomistic molecular dynamics simulations on DNA minicircles, we reveal, for the first time, the reciprocal influence between a DNA-bending protein and supercoiling. While the increased curvature of supercoiled DNA enhances wrapping around IHF, the protein pins the position of plectonemes, organizing the topology of the loop in a unique and specific manner. In addition, IHF restrains under- or overtwisted DNA depending on whether the complex is formed in negatively or positively supercoiled DNA. This effectively enables IHF to become a ‘supercoiling buffer’ that dampens changes in the surrounding superhelical stress through DNA breathing around the protein or complex dissociation. We finally provide evidence of DNA bridging by IHF and reveal that these bridges divide DNA into independent topological domains. We anticipate that the crosstalk detected here between the ‘active’ DNA and the multifaceted IHF could be common to other DNA-protein complexes relying on the deformation of DNA.

## Introduction

The recognition of specific DNA sequences by proteins is not always driven by the complementary pattern of hydrogen bonds between bases and aminoacids (so-called base or direct readout), but also can be driven by sequence-dependent deformability or local DNA structural features (indirect or shape readout) [1]. In the second mechanism, DNA is distorted in conformations that significantly deviate from the ideal B-form double helix in order to optimize the protein–DNA interface [2, 3]. Prominent examples are nucleosomes in eukaryotes and nucleic-associated proteins (NAPs) in prokaryotes, which, by bending and wrapping DNA, induce looping and other complex long-range 3D arrangements [4, 5, 6]. These DNA-bending proteins have crucial roles in organizing and packaging genomes as well as facilitating basic DNA transactions like transcription and replication [7, 8].

IHF is a key and representative NAP in Gram-negative bacteria such as *Escherichia coli* that induces one of the sharpest known DNA bends, with a measured angle of around 160^*°*^ [9]. The crystal structure reveals that IHF is formed by a core of *α* helices with a pair of extended *β*-ribbon arms whose tip each contains a conserved proline that intercalates between two base pairs [9]. These two intercalations stabilize strong bends 9 bp apart and facilitate wrapping of two DNA ‘arms’ around the protein body, tightened by electrostatic interactions between the phosphate backbone and cationic amino acids, resulting in a binding site with a length between 35-40 bp [9, 10] (Figure 1).

**Figure 1:**
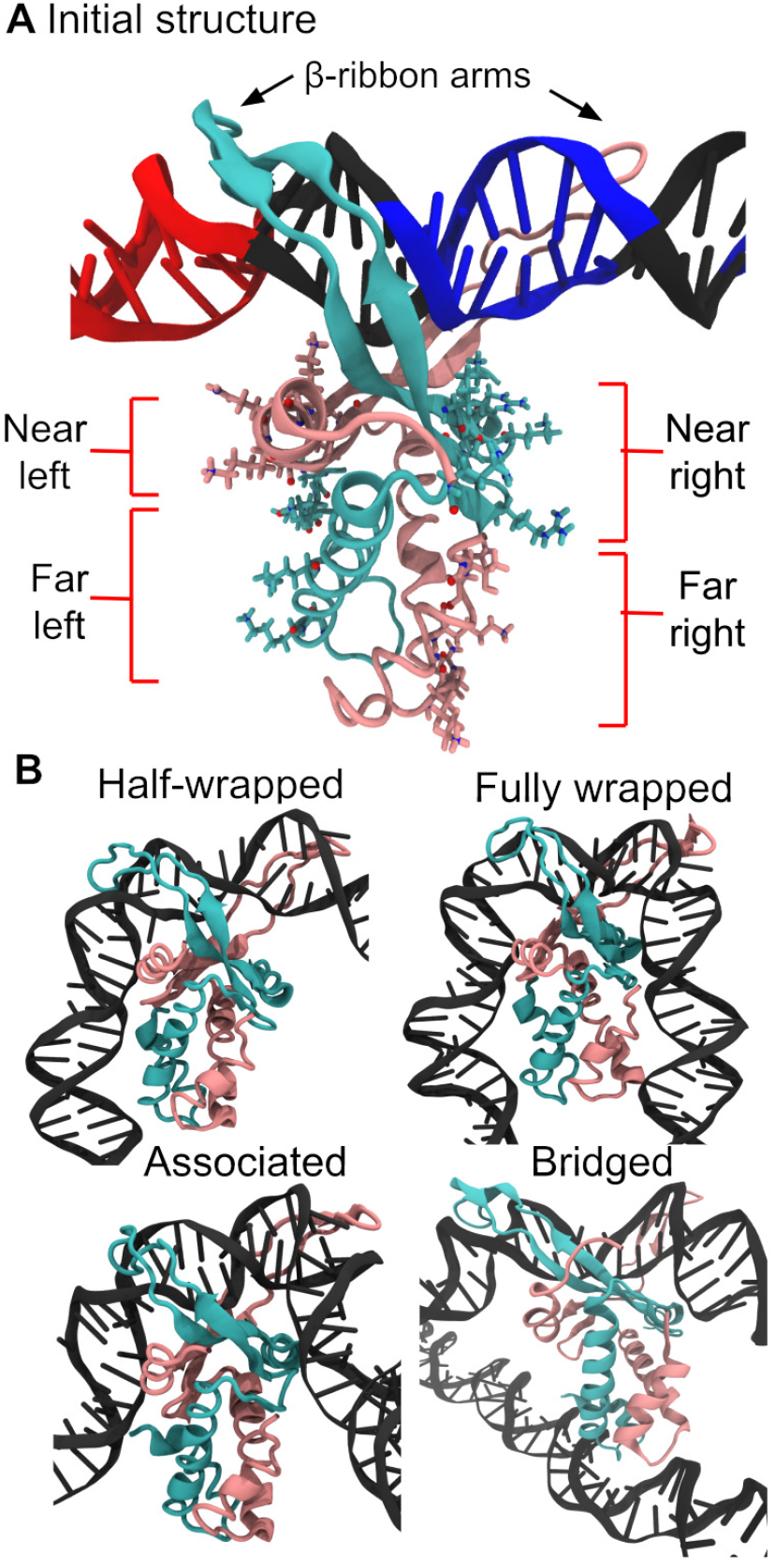
(A) Initial ‘open’ conformation for MD simulations where DNA is only bound to IHF *β*-ribon arms. (B) Linear DNA then wraps around the protein presenting two meta-stable states (half-wrapped and associated state) before arriving to the fully wrapped state, if the specific sequence is present, according to a model deduced from simulations and AFM [10]. A bridged state was also observed, in which a single copy of IHF binds to two molecules of DNA [10]. The IHF *α* subunit is shown in mauve, *β* subunit in turquoise and DNA in black except when the consensus positions are highlighted in blue and the A-tract in red. The ‘near’ and ‘far’ left sites are constituted by the *α* and *β* subunits, respectively, while the ‘near’ and ‘far’ right sites are the other way round. In the half-wrapped state, the A-tract to the left binds fully while the consensus bases to the right do not interact with the protein. In the associated state, DNA binds only to the ‘near’ sites. In the fully wrapped state, which is the one observed by crystallography, DNA arms bind to all sites

IHF binds preferentially to the DNA consensus sequence WATCARNNNNTTR (W is A or T, R is A or G, N is any nucleotide), which is located on the right side of the binding region and is small compared to the total length of the wrapped DNA [11] (Figure 1A). However, most of the strongest IHF binding sites include an A-tract to the left-hand side (upstream of the specific sequence) that increases affinity, the degree of bending and the length of the attached DNA site [12] (Figure 1A). IHF, thus, constitutes a clear example of a recognition arising through indirect readout [13, 14]. The bends induced by this protein result in higher-order structures comprising nucleoprotein complexes that are essential to a large repertoire of biological functions, including gene regulation [15], the opening of the origin of replication [16], the CRISPR-Cas system [17], and the integration and excision of phage *λ* DNA [18].

Through previous studies combining atomistic molecular dynamics (MD) simulations and atomic force microscopy (AFM), we have shown that the IHF–DNA complex is far more dynamic than previously thought [10]. Building on previous work [19], we demonstrated the existence of multiple conformations and provided structural detail of two intermediate meta-stable binding states, which are also characteristic of nonspecific DNA recognition [10]. These include a half-wrapped state in which only the upstream A-tract binds to the protein; and an associated state consisting of only partial binding on each side (see Figure 1). The fully-wrapped state, which is the one described by crystallography, is only observed in the presence of the consensus sequence, where indirect readout is induced through cooperativity between the two flanks, defining a mechanical switch on the DNA [10] (Figure 1).

We furthermore observed the formation of large DNA–IHF aggregates in AFM images and the bridging of two DNA duplexes by a single IHF protein in MD simulations (See Figure 1) [10]. This condensating behavior is of particular importance to bacterial biofilms because IHF is located at crossing points in the extracellular DNA lattice [20] and is crucial to biofilm stability [21].

In parallel, *in vivo* DNA is organized into topologically constrained domains under torsional stress [22], to which DNA responds by supercoiling. This stress causes change on the total number of DNA turns (or linking number, *Lk*) which is partitioned into twist (*Tw*) and writhe (*Wr*) as *Lk* = *Tw* + *Wr*. Structures with non-zero writhe correspond to large-scale changes in the DNA, with the helix axis twisting and bending to cross over itself, forming typically plectonemes. In the cell, DNA is maintained negatively supercoiled, with a superhelical density *σ* = Δ*Lk/Lk*_0_ ∼ − 0.06 [23, 24], being *Lk*_0_ the default linking number.

Due to inherent difficulties in obtaining high-resolution experimental structures of supercoiled DNA, computational approaches have become very useful tools [25, 26, 27], often giving excellent agreement with microscopy imaging [28, 29, 24]. In addition, computational studies have started to investigate the rich interplay between DNA topology and proteins, explaining, for instance, how the presence of proteins can shape topological domains [30, 31, 32, 5, 6]. Other studies including all-atom MD simulations on supercoiled circular DNA have found the emergence of additional secondary recognition sites between proteins and distal DNA that resulted in the formation of closed loops [33, 34]. However, to the best of our knowledge, no structural detail has been provided on the influence of torsional stress on DNA– protein interaction.

DNA supercoiling promotes the formation of its complex with IHF [35]: experiments have shown that the protein presents greater affinity for supercoiled DNA than for linear DNA [11, 36], and the disruption of the fully-wrapped state due to mutations on the lateral positions can be recovered by supercoiled DNA [37]. Of particular note is that many of the higher-order structures governed by IHF, such as integrative recombination, transcriptional regulation, and the CRISPR–Cas system, are known to be facilitated by DNA supercoiling [38, 39, 40]. Conversely, IHF has an influence on the long-range organization of DNA: the polymer is easier to circularize in the presence of the protein [36], and its knockout causes a re-organization of DNA supercoiling at the chromosome level [41].

Here, we provide atomic insight into the structural crosstalk between DNA supercoiling and protein indirect readout, using IHF as a model case of study. This protein is a remarkable example as it induces one of the sharpest bends on DNA. By simulating the dynamics of DNA minicircles bound to IHF, we identify the importance of supercoiling to the protein’s binding mode when relying on indirect readout. We observe that enhancement on DNA flexibility and curvature by supercoiling leads to an increase of DNA-binding modes with a tendency to enhance wrapping around the protein. We also explore the entropic reduction of the conformational landscape of supercoiled DNA by IHF, as well as its capacity to constrain superhelical stress. We finally provide further insight into the formation of closed DNA loops bridged by IHF and demonstrate the formation of independent topological domains.

## MATERIALS AND METHODS

### Construction of DNA minicircles

A linear 336 bp DNA fragment was built using the NAB module implemented in Amber16 [42] with a sequence based on the minicircle generated by intramolecular *λ*-integrase recombination [43, 29]. This sequence, containing a single IHF binding site, is given in Section 1 of the supplementary material. Six perfectly planar DNA minicircles containing between 29 and 34 turns were then constructed using an in-house program as previously performed [24]. Afterwards, the structure of IHF-DNA from phage *λ* excision complex (Protein Data Bank (PDB): 5J0N [18]) was inserted at the matching IHF-binding H2 site contained at the attR region of the minicircle. Only the central 11-bp from H2 site that enclose the two intercalation sites was replaced by the crystallographic structure and then junctions between DNA fragments were minimized until a canonical structure was achieved, following previous studies [34]. Hence, the resultant structure used to start simulations consisted of DNA minicircles bound to IHF in an ‘open state’ without lateral interactions (see Figure 1).

### Molecular dynamics simulations

All simulations were set up with the AMBER 16 suite of programs and performed using the CUDA implementation of AMBER’s pmemd program [42]. The constructs were solvated using implicit generalized Born model at a sodium chloride salt concentration of 0.2 M with GBneck2 corrections, mbondi3 Born radii set and no cut-off for a better reproduction of molecular surfaces, salt bridges and solvation forces [44, 45, 46]. Langevin dynamics was employed for temperature regulation at 300 K with a collision frequency of 0.01 ps^1^ in order to reduce the effective solvent viscosity and, thus, accelerate the exploration of conformational space [47, 10]. The protein and DNA were represented by ff14SB [48] and BSC1 [49] force fields, respectively. Prolines were kept intercalated by restraining the distances between key atoms in the proline side chain and neighboring bases [10]. Following our protocols for minimization and equilibration [10], three replica simulations of 30 ns each were performed for each topoisomer with IHF bound, and three more for the same systems with the protein removed. The first 20 ns were obtained with distance restraints on the WC canonical H-bonds to avoid a premature disruption of the double helix [34], so only the last 10 ns of each simulation were considered for analysis.

### Analysis of simulations

Topological DNA twist and writhe were calculated using WrLINE, which outputs global twist and writhe values alongside the molecular contour [50]. Because global and local definitions of twist are not directly compatible [51], the accumulative twist at the DNA binding site was calculated according to the 3DNA definition at the dinucleotide level [52] using SerraNA [53]. Simulations in implicit solvent are known to systematically overestimate DNA twist [54]. To correct this, a linear fit of average writhe for bare minicircles was performed, so we could determine the value of *Lk* for which *Wr* = 0 (Figure S1); this was found to be *Lk*_0_ = 31.08. Then, *σ* for each topoisomer was calculated relative to this value.

Hydrogen bonds were determined using cpptraj [55] with a distance cutoff of 3.5 Å and an angle cut-off of 120^*°*^. The number of hydrogen bonds involving each protein residue and DNA was capped at one, so time-averages along trajectories indicate the proportion of frames presenting this interaction. This was compared with the hydrogen bonds presented in the original crystallographic structure, which is the PDB entry 1IHF [9]. It should be noted that PDB 5J0N was obtained via CryoEM and posterior fitting based on 1IHF.

All simulation frames were classified via hierarchical agglomerative clustering based on the average linkage algorithm using root-mean-squared deviation (RMSd) between frames as a distance metric [55]. Only the backbone atoms of IHF and of a 61 bp region of DNA centered on the binding site were considered for the RMSd. The number of clusters was chosen so each had a distinct interaction pattern of hydrogen bonds between the protein and DNA.

## RESULTS AND DISCUSSION

Six different topoisomers (ΔLk=-2,-1,0,1,2,3) of DNA minircircles containing 336 bp were constructed in order to achieve a similar *σ* range to the one observed *in vivo* (from -0.067 to +0.094). Then, these were attached to IHF via only its protruding *β*-ribbon arms to simulate how DNA spontaneously wraps around the protein following an initial bound state, which resembles an encounter complex formed at the beginning of the recognition process (Figure 1) [56, 14, 10].

Three independent MD simulation replicas were performed for each topoisomer with/without IHF in implicit solvent to allow enough conformational sampling over feasible timescales (see Supplementary Movies 1-12). A continuum representation of the solvent reduces the computational cost of simulations compared with a solvation box with discrete water molecules and ions, and accelerates global structural rearrangements by at least an order of magnitude due to the neglect of solvent viscosity [29]. Although non-bonded interactions are not so accurately described on implicit solvent, we found these type of simulations reproduced well X-ray IHF-DNA interactions and the induced DNA bends measured by AFM on linear DNA [10]

### DNA conformation has an active role in indirect-readout recognition

To identify the principal DNA-binding modes, all frames from all trajectories were merged together and classified into five distinct binding modes (Figure 2A) presenting a characteristic DNA-protein interaction pattern (Figure 2B) (see Methods).

**Figure 2:**
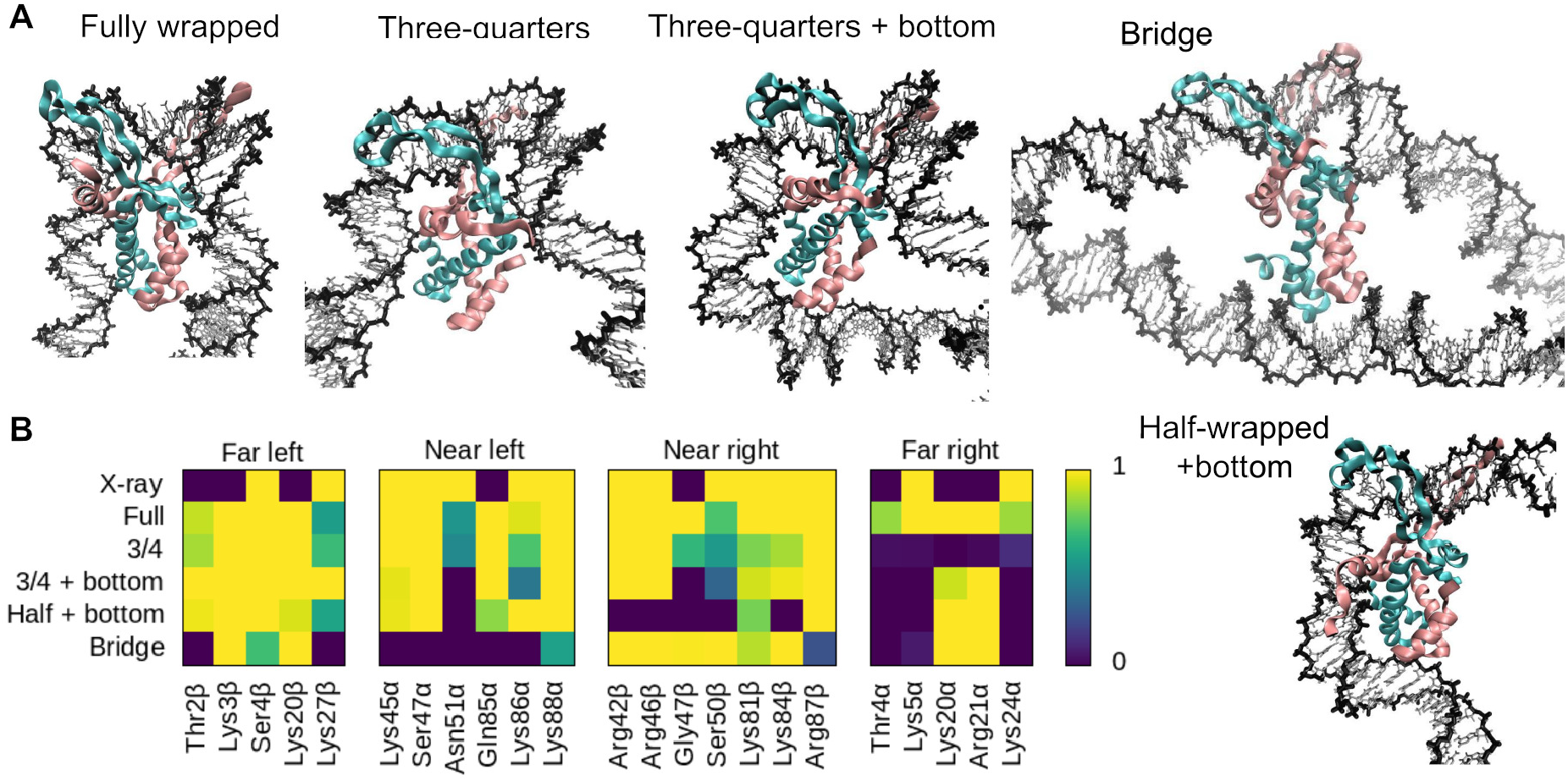
**(A)** Representative structures of the different binding modes observed in our simulations. **(B)** Time-average number of intermolecular hydrogen bonds formed by the main DNA-interacting amino acids belonging to each binding mode, with the crystal structure (PDB 1IHF; labeled X-ray) provided for comparison; note that the DNA in this structure is too short (35 bp) to capture some interactions in the ‘far’ regions. Color scheme is the same as in Figure 1.

As has been described previously [10], interactions between IHF and the lateral DNA arms can be divided into four regions based on their position relative to the center of the binding site and the protein subunit to which the involved amino acid belongs. On the left-hand side (containing the A-tract), the *α* subunit is closer to the center and thus constitutes the ‘near left’ site, while the *β* subunit is farther and composes the ‘far left’. On the right-hand side (containing the consensus sequence), the *α* and *β* subunits are inversely arranged, delimiting the ‘far right’ and ‘near right’ sites, respectively (see Figure 1A).

As expected, the fully wrapped state is observed, presenting very similar protein-DNA contacts to the crystal structure [9] (Figure 2). The half-wrapped and associated states previously observed for linear DNA (Figure 1) do not appear, probably due to the inherent curvature of circular DNA (around 64^*°*^ over a region the length of the IHF-interacting site), which can be expected to bias the system towards more tightly wrapped states. Instead, a ‘three-quarters’ state emerges in which the A-tract on the left binds fully to the protein while the right DNA arm binds only to the near right site. Two extra new states appear, both involving the binding of the left DNA arm to the “bottom” of the protein, while the right arm remains either unbound (‘half-wrapped + bottom’) or bound to only the near site (‘three-quarters + bottom’) (see Figure 2). Lysine 20 and Arginine 21 from the subunit *α* at the far right site are the aminoacids mainly responsible to wrap the left DNA arm around the “bottom” of the protein (Figure 2). We also observed state comprising an IHF-mediated DNA bridge similar to those previously demonstrated [10], where the DNA remains relatively unbent and the two far sites or the “bottom” of the protein interact with a second DNA double helix (Figure 2).

We barely observe transitions between states over time within individual replica simulations (see Supplementary Movies 1-12). As Table 1 shows, only one simulation is observed to sample several conformations: replica 1 for the most relaxed topoisomer switches from the three-quarters to the fully wrapped state (Supplementary Movie 5). This suggests that all of these observed binding modes are stable states corresponding to free-energy minima, where the simulations are trapped, rather than temporary transition structures en route to a global minimum.

**Table 1:**
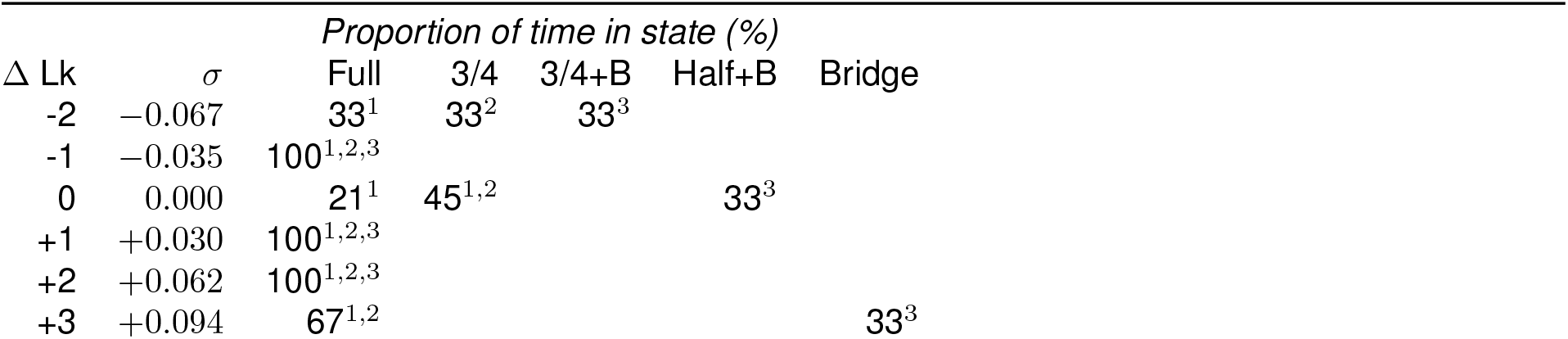
Populations of the conformational states vary with the superhelical density of DNA. Percentage of simulation frames for each state and topoisomer, where ‘Full’ refers to the fully wrapped state, ‘3/4’ to the three-quarters, ‘3/4+B’ to three-quarters + bottom, ‘Half+B’ to half-wrapped + bottom and ‘Bridge’ to IHF-mediated DNA bridge

Because the final complex interactions are determined by the intrinsic structure and dynamics of DNA, our simulations demonstrate that DNA is not just a passive polymer to be manipulated, but it has an active role in driving the IHF recognition process [35].

### Supercoiling affects DNA recognition by IHF

We find that the populations of these states vary with the superhelical density of DNA (Table 1 and Figure 3). While relaxed minicircles present the fully wrapped state, they show a preference for more open states like three-quarters and half-wrapped + bottom (Supplementary Movies 5-7). The propensity for the fully wrapped state is strongly enhanced for moderate levels of positive and negative supercoiling, as this binding mode is presented exclusively for topoisomers ΔLk=-1,1 and 2 (Supplementary Movies 4,8 and 9). Hence, our simulations reveal that an increase in the underlying DNA curvature induced by supercoiling significantly facilitates DNA-shape readout by IHF, promoting larger wrapping around the protein compared with relaxed DNA.

**Figure 3:**
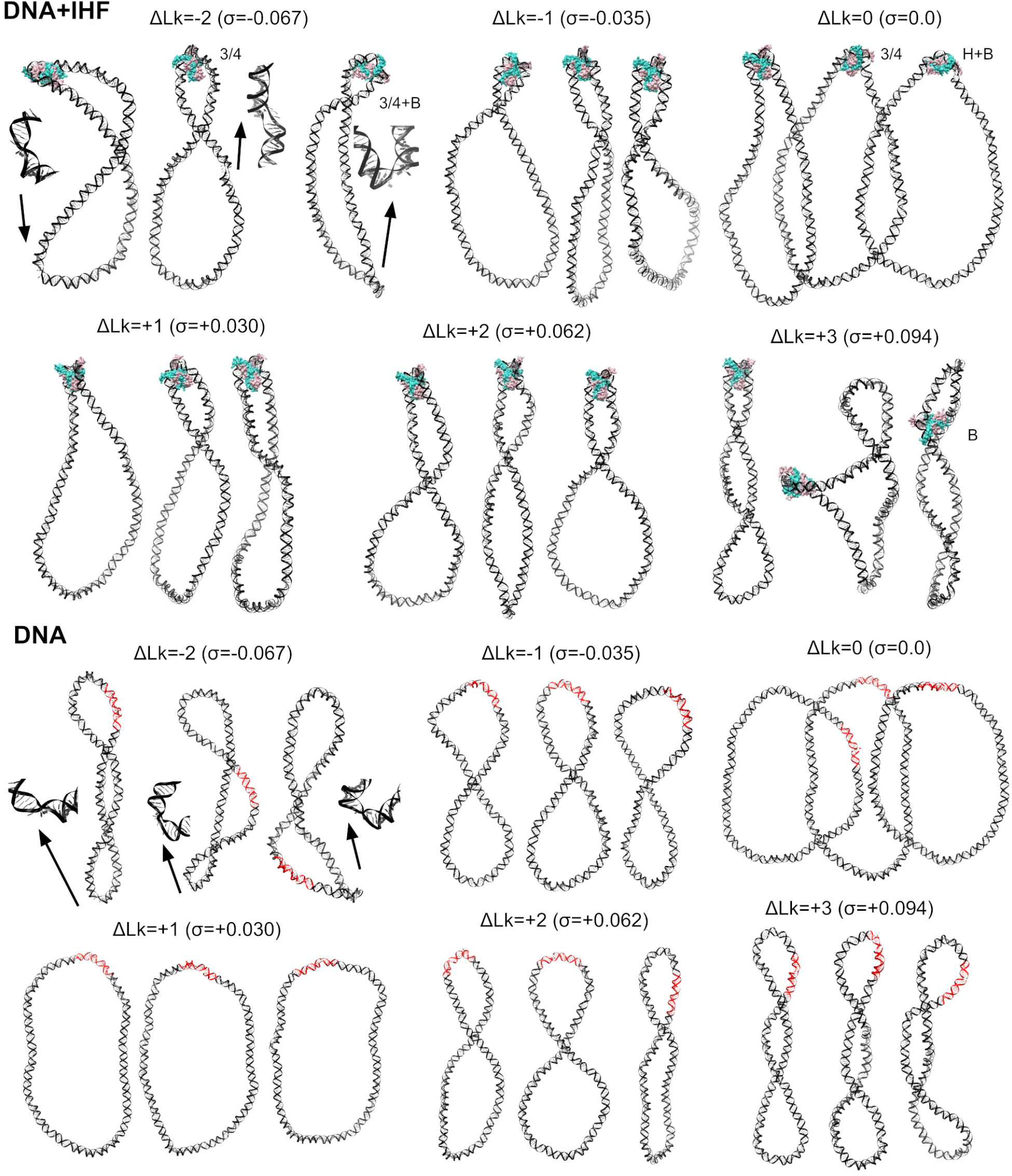
Overview of the dependence of the DNA-IHF interaction landscape on superhelical density given by representative structures for each individual simulation. Replicas 1, 2 and 3 for each topoisomer are displayed from left to right, respectively, and labeled with the binding mode when they not present the fully wrapped: ‘3/4’ for the three-quarters, ‘3/4+B’ for three-quarters + bottom, ‘H+B’ to halfwrapped + bottom and ‘B’ to IHF-mediated DNA bridge. These extra states are mainly presented on relaxed DNA (ΔLk=0) for more open conformations and on highly supercoiled DNA (ΔLk=-2,+3) promoted by enhanced flexibility and defects on the double helix (zoom-ins, indicated by arrows). IHF is mostly located at the apex of plectonemes, as opposed to bare DNA where IHF binding sites (in red) have multiple locations. Color scheme is the same as in Figure 1.

We find that readout variability increases for higher superhelical densities (Figure 3): the most negatively supercoiled topoisomer (ΔLk=-2) presents different binding modes per each replica (see Supplementary Movies 1-3); the most positively supercoiled topoisomer (ΔLk=+3) results in a compact trefoil conformation in its second replica (Supplementary Movie 11) and an IHF-mediated bridge in its third (Supplementary Movie 12). As the level of torsional stress increases, DNA tends to present a broader distribution of conformations due to the emergence of extra supercoiled bends and defects in the double helix [57, 24]. These defects are associated with a wider ensemble of possible structures, because they occur stochastically at multiple sites [58, 57] and act as flexible hinges, allowing stress release and significant structural readjustments [34]. We observe the emergence of denaturation bubbles in all replica simulations of topoisomer ΔLk=-2 (see Figure 3), which presents a superhelical density close to that steadily maintained in most live bacteria (*σ*=-0.067) [23, 24]. This, and the fact that the extent of supercoiling widely differs between chromosomal regions [59], make the observed variability likely to be present *in vivo*.

### The effect of IHF on minicircle compactness and twist-writhe partition

Our simulations show that IHF globally compacts relaxed DNA loops and that this effect is proportional to the level of wrapping around the protein (Figure 4). The first replica of topoisomer ΔLk=0, where DNA is fully wrapped, presents the strongest reduction in the radius of gyration compared with the second replica, where the DNA is wrapped three-quarter parts, and the third, where the DNA is only half wrapped (Figure 4).

**Figure 4:**
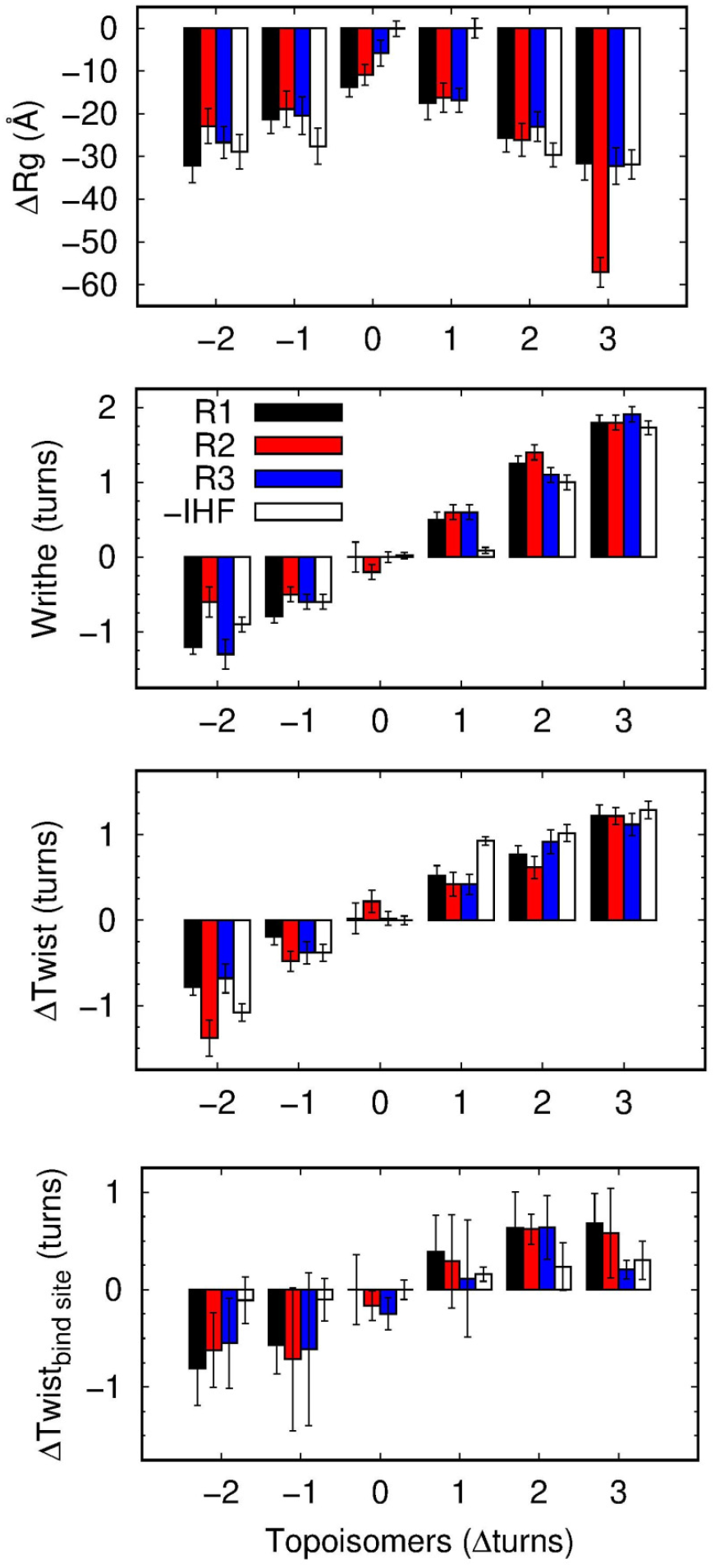
Averages and corresponding standard deviation (error bars) of radius of gyration (a), writhe (b), twist for the whole circle (c) and twist on the IHF binding site (d) of DNA minicircles with different levels of supercoiling, with IHF (black, red and blue for replicas 1, 2 and 3, respectively) and without IHF (white). Replica simulations are ordered from left to right as in Figure 3. The extremely low value in the radius of gyration observed for the 2nd replica of ΔLk=+3 is due to the formation of a highly compact trefoil structure (see Figure 3).

As the degree of supercoiling increases in either direction, this compaction effect becomes superfluous, as DNA naturally becomes rod-like (see Figure 3 and 4). An exception to this is the ΔLk=+1 topoisomer, which remains predominantly open in the absence of IHF and becomes substantially compacted upon protein binding (Figure 4). IHF also brings a significant change in the twist-writhe partition on this topoisomer, which has the effect of correcting the asymmetry between positively and negatively supercoiled DNA (see Figure 4). On naked DNA, negative supercoiling is associated with more writhed structures than equivalent amounts of positive supercoiling (Figure 3). However, IHF appears to correct this asymmetry by shifting the writhe of ΔLk=+1 topoisomer in the positive direction. Because twist at the binding site cannot explain the altered twist-writhe balance (Figure 4), we hypothesize that this effect is due to IHF-mediated bends, which stimulate writhed apex-like structures (Figure 3), enabling twist relaxation.

Finally, we relate twist-writhe variability observed in topoisomer ΔLk=-2 to the presence of DNA defects (Figure 4). Replica 2 presents a bigger denaturation bubble compared with the other two replicas (Figure 3), which causes extremely low twist values and, as a result, a considerable moderation in writhe (Figure 4).

### IHF restrains under- or overtwisted DNA depending on supercoiling direction

Figure 4 shows that DNA twist at the binding site presents lower or higher values (between 0.5 to 1 helical turn) compared with relaxed DNA, depending on whether the complex is formed under negatively or positively supercoiled DNA, respectively. To understand the origin of this effect, we looked into the structures in detail and we observed a considerable amount of heterogeneity as DNA is wrapped around the protein under different levels of torsional stress (Figure 5 and S2). These conformational adjustments, which mainly consist of changes in molecular twist and groove dimensions, induce the protein to interact with different nucleotides, thus pinning the double helix in distinct orientations (Figure 5 and S2).

**Figure 5:**
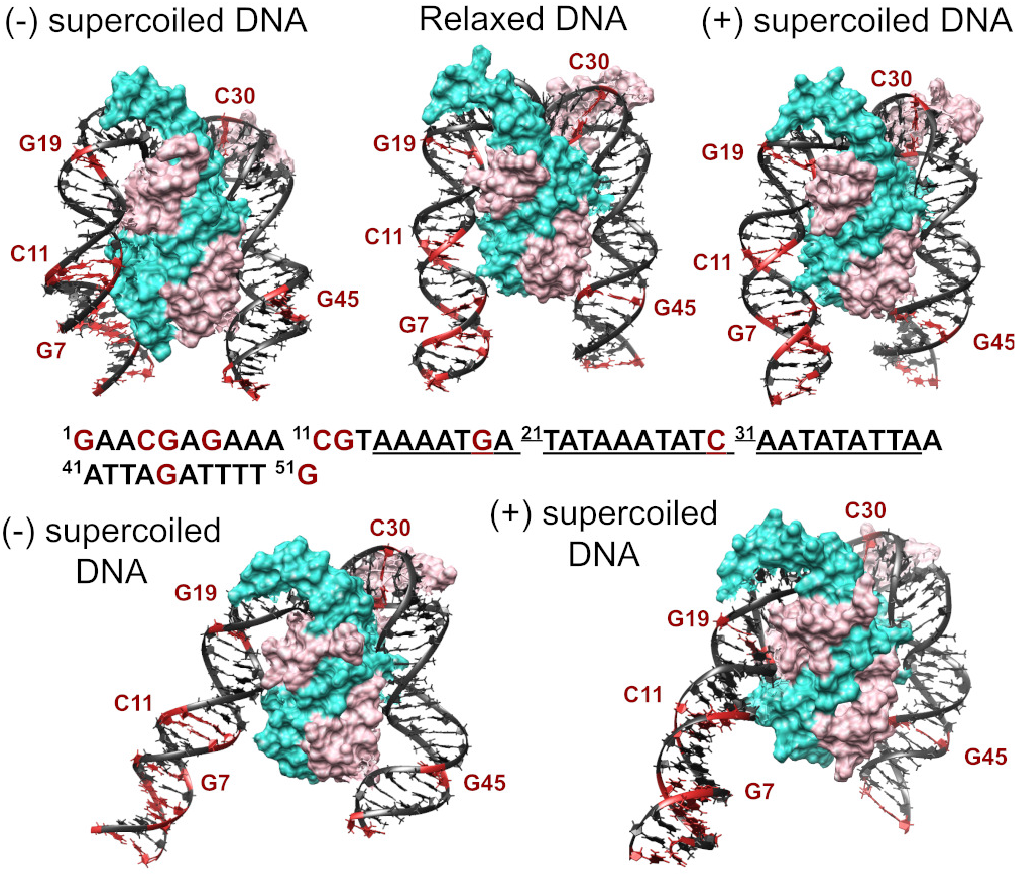
Supercoiling dependence of DNA structure when bound to IHF. Representative examples of the fully wrapped state formed under negatively (left) and positively (right) supercoiling reveal changes in twist compared with relaxed DNA (middle) (see Figure S2 for all replicas). The complete DNA sequence is included, where the consensus binding site is in underlined text and the most conserved positions in bold. The only few CG bp are highlighted in red and serve as rulers to compare DNA orientation relative to IHF sides. The rest of the color scheme is the same as in Figure 1. The two bottom structures reveal variability in the supercoiled DNA being fully wrapped around the protein with sizable changes in groove dimensions (right side) and a reduction in the contact points (left side).

We also find that, on occasion, DNA supercoiling reduces the number of contact points between a DNA arm and its IHF side from three (encompassing two major and one minor grooves) to two (a major and a minor groove) (see the two bottom structures of Figure 5). We do not observe this conformational alteration in relaxed DNA, probably due to its natural propensity to optimally wrap IHF. Hence, our simulations reveal that the DNA conformational variability induced by supercoiling not only influences the binding modes of the complex but also its fine structural details.

Previous experiments have given an unclear picture of whether IHF constrains supercoiled DNA: while *in vivo* experiments found IHF was not able to change the overall supercoiling balance in the chromosome [60, 59], *in vitro* experiments showed that IHF had indeed the capacity to constrain supercoiled DNA on smaller plasmids [36]. Our simulations provide an explanation for these apparently contradictory results: IHF can restrain twist at the binding site, although it cannot modify the global state, because it under- or overwinds DNA depending on the supercoiling direction. In fact, our results suggest that IHF could act as a kind of ‘supercoiling buffer’ through the release of stored torsional stress by means of DNA breathing or dissociation as the surrounding superhelical density would change. This view is in overall agreement with a recent study reporting redistribution of DNA supercoils at the chromosome level when IHF is eliminated [41].

### IHF reduces the entropy of the DNA supercoiling conformational landscape

In the presence of IHF, plectonemes are mostly observed to form with the protein at their apices (see Figure 3 and 6). This has the effect of significantly reducing the entropy of the minicircle conformational landscape, relative to the case in which no protein is bound (Figure 6). We observe that the conformational distribution of the DNA minicircles is significantly broader in naked DNA, as the apex of the plectoneme can be located in multiple positions. In the presence of the protein, the ensemble of conformational states is shifted towards a unique folded state, positioning the IHF at the apex.

**Figure 6:**
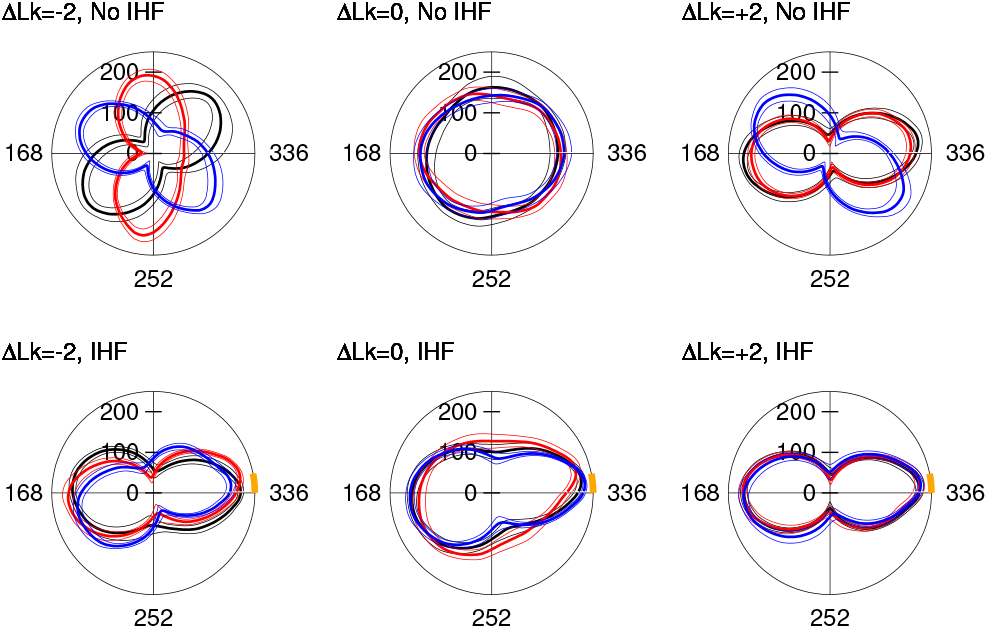
Polar plots of mean (± standard deviation) distance from each point along the helix axis to the DNA centroid. Each color represents a replica simulation. In the absence of IHF (top), plectonemes can form in many positions. Adding IHF (bottom, binding site location shown in orange) causes the plectonemes to align with the protein at the apex and consistently localize to the crossing points.

We can quantitatively estimate the cost of the entropy reduction by using *S* = *k*_*B*_*ln*(*W*), where *k*_*B*_ is the Boltzmann constant and *W* is the number of possible states. If we assume IHF folds DNA in one state, compared with the 168 possible in naked DNA (an apex of the plectoneme can be pinned to each bp along half of the minicircle), then the entropy reduction is approximately 5.1 *k*_*B*_*T* or 3 kcal/mol at 300 K. If we consider that not all plectoneme positions are equally probable along the naked minicircle (some conformations are more favorable than others, see Figure 3 and 6), we then need to reduce the number of states to 50 or 25%. This gives entropy penalties around 4.4 *k*_*B*_*T* (2.6 kcal/mol) and 3.7 *k*_*B*_*T* (2.2 kcal/mol), respectively, which are still large enough to be overcome by thermal fluctuations of bare DNA. This entropic simplification could be larger, as IHF could have the capacity to organize longer DNA loops, containing higher levels of inherent conformational variability.

Interestingly, a similar plectoneme-pinning effect has also been detected in damaged DNA [61, 62], showing that local changes in DNA curvature and flexibility are key to regulating the folding of supercoiled loops. Overall, our simulations support the view that the mechanism of IHF action is to organize DNA into unique conformations in order to facilitate all types of genetic transactions in which the protein is involved.

### IHF-mediated bridging divides DNA into topological domains

A DNA–IHF–DNA bridge involving additional contacts between distal DNA and the “bottom” of the protein was observed to form spontaneously in replica 3 of the most positively supercoiled minicircle (Δ*Lk* = +3) (see Supplementary Movie 12). This bridge results from nonspecific interactions between basic aminoacids and the negatively charged DNA backbone (see Figure 2 and 3). This supports our previous findings indicating that such bridges are both possible and energetically favorable, and that specific recognition can be simply modulated or extended via additional electrostatic-driven interactions between the protein and the DNA [10].

The observation of this bridge in the most supercoiled minicircle can be explained by the proximity of distal DNA sites that are far apart in torsionally relaxed DNA, as has been suggested by single-molecule experiments [39, 63, 64]. In this regard, DNA bridges involving secondary nonspecific recognition sites have also been identified for other bacterial proteins like Topoisomerase IB [33] and ParB [65] in super-coiled DNA.

The formation of an IHF-mediated DNA bridge in a minicircle results in two closed loops. Measuring the writhe in both of these loops over time (Figure 7), reveals no evidence of writhe passing between the loops, consistent with the formation of two isolated topological domains. Furthermore, the writhe is not evenly distributed: while the larger loop accounts for 76% of the minicircle’s contour length (255 bp), it holds 90% of the total writhe. That this asymmetry was not corrected by the diffusion of writhe into the smaller loop is further evidence for the separation of topological domains.

**Figure 7:**
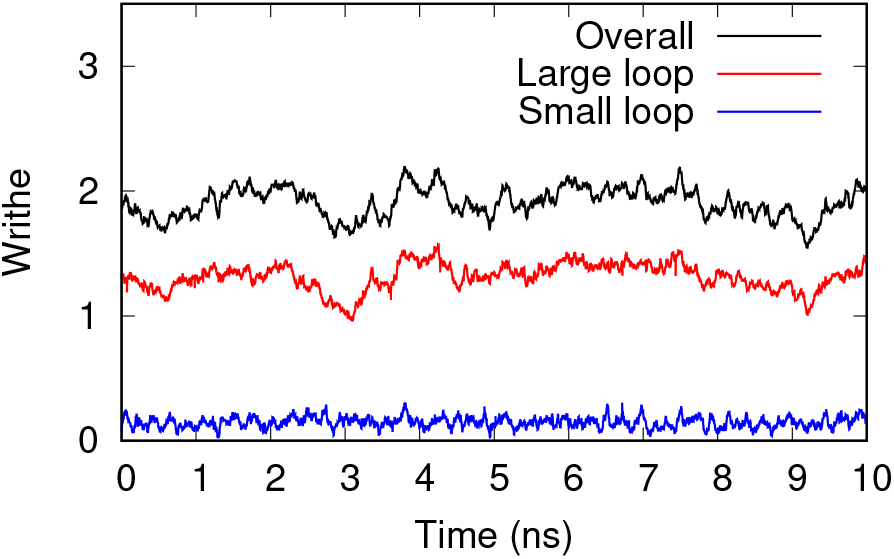
A DNA minicircle bridged by IHF consists of two closed loops, each of which has independent writhe. There is no evidence of writhe passing between them: the writhe of the smaller loop (81 bp, blue) remains constant, while any changes in the writhe of the larger loop (255 bp, red) are also reflected in the minicircle’s overall writhe (336 bp, black).

This effect can be quantified by calculating the correlation coefficients between each pair of time series: if writhe regularly passes between the two loops, one would expect the two datasets to be negatively correlated with *R*^2^ close to 1. In fact, the calculated value is *R*^2^ = 0.0041, indicating that no correlation exists between the two and that IHF is therefore demonstrably dividing the DNA minicircle into two separate topological domains. For comparison, the *R*^2^ values for the correlation of the overall writhe with the large and small loops are 0.75 and 0.14, respectively, indicating as expected that the larger loop has a greater influence on the total writhe and that changes within both loops collectively explain almost all of the change in the minicircle’s overall writhe.

Finzi and coworkers have already shown that protein-mediated DNA bridges have the capacity to establish independent topological domains, constraining variable amounts of supercoiling [66, 67]. This result was observed by specialized loop-mediating proteins like the CI [66] and *lac* repressors [67], where each DNA molecule is attached to the bridging protein by means of specific interactions. Here, our simulations provide atomic insight into this effect and reveal that a single bridge is sufficient to create a topological boundary, even if it is locked via nonspecific interactions.

## CONCLUSIONS

By performing all-atom simulations, we have provided, for the first time, atom-level insights of the interplay between DNA supercoiling and DNA-shape protein recognition (see Figure 8). We observe that changes in the intrinsic curvature of circular DNA facilitates its bending around IHF and results in the appearance of new binding modes not observed in relaxed linear DNA [10]. We also observe that these effects are further enhanced by supercoiled DNA (Figure 8A). Because this study examines DNA supercoiling within ranges observed *in vivo*, we expect our findings to be relevant in the living cell. We anticipate that the DNA’s ‘active role’ [35] detected here will be applicable to other systems relying on indirect recognition, where DNA is heavily deformed, including other NAPs and eukaryotic chromatin-binding proteins.

**Figure 8:**
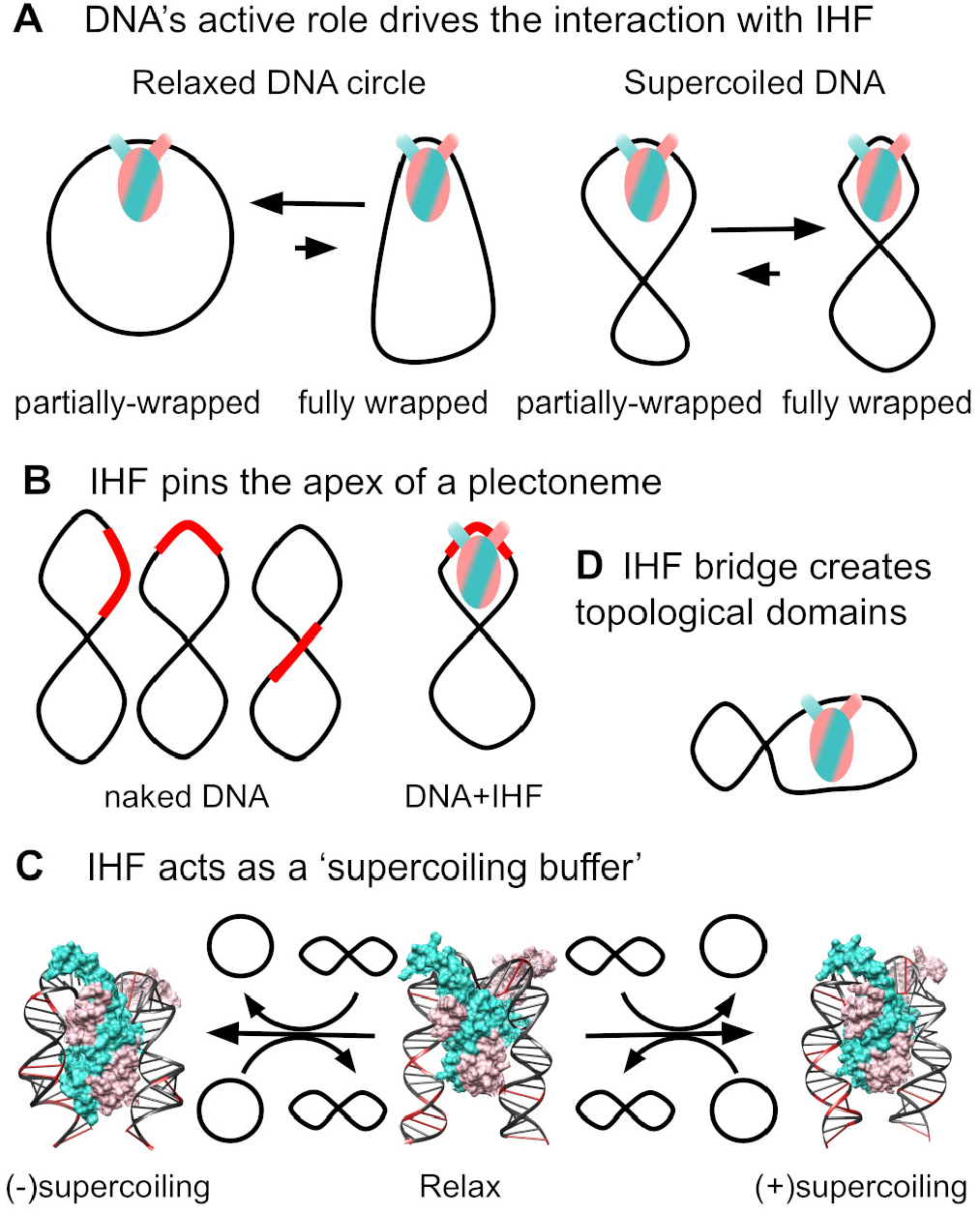
Model of the different ways through which the interplay between a DNA-bending protein like IHF and supercoiling emerges. (**A**) DNA intrinsic curvature facilitates wrapping around the protein, thus demonstrating DNA’s active role in this recognition. (**B**) In reverse, IHF organizes DNA conformation in a unique manner through the pinning of plectoneme’s apex. In bare DNA, the binding site (in red) can be in a variety of positions. (**C**) IHF can also act as a ‘supercoiling buffer’ by restraining underor overtwisted DNA at the binding site depending on whether the complex is formed on negatively or positively supercoiled DNA. As the neighboring supercoiling decreases or increases (represented by an open circle or an ‘8’ shape, respectively), the constrained twisting is released, shielding a steady supercoiling level. CG bps are in red and serve as rulers to identify differences in DNA helical pitch. (**D**) Finally, IHF can separate DNA in topological domains through a non-specific electrostatic-driven bridging interaction. The color scheme is the same as in Figure 5.

As well as quantifying the influence of supercoiling on IHF binding, we also demonstrate the effect of IHF binding on the topological organization of DNA by showing that IHF strongly and reliably controls the position of plectonemes (Figure 8B). We have shown that funneling the DNA energy landscape towards one particular topological arrangement is not trivial and that this is a key mechanism through which IHF exerts its biological functions.

We have also detected the capacity of IHF to restrain under- or overtwisted DNA in its binding site depending on whether the complex is formed under negatively or positively supercoiled DNA. This effect suggests that the protein could act as a ‘supercoiling buffer’ by increasing or decreasing constrained supercoiled DNA as neighboring superhelical density is changed (Figure 8C) via DNA breathing or complex dissociation, thus, shielding a steady state of supercoiling.

Additional evidence [10] is also provided for DNA bridging by IHF when a second double helix is nearby, via a secondary nonspecific interaction (Figure 8D). Probably, this is of significance to a number of biofilms and to nucleoid compaction at the cellular stage when IHF is most abundant. We demonstrate that this bridging, even if it is based on nonspecific interactions, has the capacity to divide the DNA into two distinct topological domains as it prevents the transfer of supercoiling between them.

The combination of these effects provides a biological mechanism to control plectoneme positions, supercoiling and chromosome boundaries, making IHF a valuable tool for the regulation of genes in complex pathways as has been detected at the whole genomic level [41]. We anticipate that this multifaceted mode of action might not be exclusive of IHF, but it could constitute a common principle of architectural proteins responsible for the organization of chromosomes, either in prokaryotes or eukaryotes, and, more generally, of proteins that recognises DNA through alterations on its shape.

## ACKNOWLEDGEMENTS

Engineering and Physical Sciences Research Council (EPSRC) [EP/N027639/1, EP/T002166/1, EP/R029407/1, EP/P020259/1]; Biology and Biotechnology Research Council (BBSRC) [BB/R001235/1]; Leverhulme Trust [RPG-2017-340]; University of York Research Priming funds [ref 50109436 and G0081301]. Calculations were performed on ARCHER, JADE, Cambridge Tier-2 and the local York facilities (Viking and YARCC clusters). Funding for open access charge: York Open Access Fund. We would like to thank Lynn Zechiedrich, Jonathan M. Fogg, Tom C. B. McLeish and Sarah A. Harris for useful discussions and comments.

## References

[1] Rohs, R., West, S. M., Sosinsky, A., Liu, P., Mann, R. S., and Honig, B. (2009) The role of DNA shape in protein–DNA recognition. Nature, 461, 1248–1253.

[2] Li, J., Sagendorf, J. M., Chiu, T.-P., Pasi, M., Perez, A., and Rohs, R. (2017) Expanding the repertoire of DNA shape features for genome-scale studies of transcription factor binding. Nucleic Acids Res., 45, 12877–12887.

[3] Battistini, F., Hospital, A., Buitrago, D., Gallego, D., Dans, P. D., Gelpí, J. L., and Orozco, M. (2019) How B-DNA Dynamics Decipher Sequence-Selective Protein Recognition. J. Mol. Biol., 431, 3845–3859.

[4] Collepardo-Guevara, R. and Schlick, T. (2014) Chromatin fiber polymorphism triggered by variations of DNA linker lengths. Proc. Natl. Acad. Sci. U.S.A., 111, 8061–8066.

[5] Todolli, S., Young, R. T., Watkins, A. S., Bu Sha, A., Yager, J., and Olson, W. K. (2021) Surprising Twists in Nucleosomal DNA with Implication for Higher-order Folding. J. Mol. Biol., 433, 167121.

[6] Clauvelin, N. and Olson, W. K. (2021) Synergy between Protein Positioning and DNA Elasticity: Energy Minimization of Protein-Decorated DNA Minicircles. J. Phys. Chem. B, 125, 2277–2287.

[7] Bai, L. and Morozov, A. V. (2010) Gene regulation by nucleosome positioning. Trends in Genetics, 26, 476–483.

[8] Dame, R. T., Rashid, F.-Z. M., and Grainger, D. C. (2020) Chromosome organization in bacteria: mechanistic insights into genome structure and function. Nat. Rev Genet., 21, 226–242.

[9] Rice, P. A., Yang, S. W., Mizuuchi, K., and Nash, H. A. (1996) Crystal structure of an IHF-DNA complex: A protein-induced DNA U-turn. Cell, 87, 1295–1306.

[10] Yoshua, S., Watson, G., Howard, J. A. L., Velasco-Berrelleza, V., Leake, M., and Noy, A. (2021) Integration host factor bends and bridges DNA in a multiplicity of binding modes with varying specificity. Nucleic Acids Res., 49, 8684–8698.

[11] Yang, S. W. and Nash, H. A. (1995) Comparison of protein binding to DNA in vivo and in vitro: defining an effective intracellular target.. EMBO J., 14, 6292–6300.

[12] Hales, L. M., Gumport, R. I., and Gardner, J. F. (1996) Examining the contribution of a dA+dT element to the conformation of Escherichia coli integration host factor-DNA complexes. Nucleic Acids Res., 24, 1780–1786.

[13] Swinger, K. K. and Rice, P. A. (2004) Ihf and hu: flexible architects of bent dna. Curr. Opin. Struct. Biol., 14, 28–35.

[14] Velmurugu, Y., Vivas, P., Connolly, M., Kuznetsov, S. V., Rice, P. A., and Ansari, A. (2018) Two-step interrogation then recognition of DNA binding site by Integration Host Factor: An architectural DNA-bending protein. Nucleic Acids Res., 46, 1741–1755.

[15] Huo, Y. X., Zhang, Y. T., Xiao, Y., Zhang, X., Buck, M., Kolb, A., and Wang, Y. P. (2009) IHF-binding sites inhibit DNA loop formation and transcription initiation. Nucleic Acids Res., 37, 3878–3886.

[16] Hwang, D. and Kornberg, A. (1992) Opening of the replication origin of Escherichia coli by DnaA protein with protein HU or IHF. J. Biol. Chem., 267, 23083–23086.

[17] Wright, A. V., Liu, J.-J., Knott, G. J., Doxzen, K. W., Nogales, E., and Doudna, J. A. (2017) Structures of the CRISPR genome integration complex. Science, 357, 1113–1118.

[18] Laxmikanthan, G., Xu, C., Brilot, A. F., Warren, D., Steele, L., Seah, N., Tong, W., Grigorieff, N., Landy, A., and Van Duyne, G. D. (2016) Structure of a holliday junction complex reveals mechanisms governing a highly regulated DNA transaction. eLife, 5, 1–23.

[19] Connolly, M., Arra, A., Zvoda, V., Steinbach, P. J., Rice, P. A., and Ansari, A. (2018) Static Kinks or Flexible Hinges: Multiple Conformations of Bent DNA Bound to Integration Host Factor Revealed by Fluorescence Lifetime Measurements. J. Phys. Chem. B, 122, 11519–11534.

[20] Novotny, L. A., Amer, A. O., Brockson, M. E., Goodman, S. D., and Bakaletz, L. O. (2013) Structural Stability of Burkholderia cenocepacia Biofilms Is Reliant on eDNA Structure and Presence of a Bacterial Nucleic Acid Binding Protein. PLOS ONE, 8, e67629.

[21] Devaraj, A., Justice, S. S., Bakaletz, L. O., and Goodman, S. D. (2015) DNABII proteins play a central role in UPEC biofilm structure. Mol. Microbiol., 96, 1119–1135.

[22] Postow, L., Hardy, C. D., Arsuaga, J., and Cozzarelli, N. R. (2004) Topological domain structure of the Escherichia coli chromosome. Genes Develop., 18, 1766–1779.

[23] Zechiedrich, E., Khodursky, A. B., Bachellier, S., Schneider, R., Chen, D., Lilley, D. M., and Cozzarelli, N. R. (2000) Roles of Topoisomerases in Maintaining Steady-state DNA Supercoiling in Escherichia coli. J. Biol. Chem., 275, 8103–8113.

[24] Pyne, A. L. B., Noy, A., Main, K. H. S., Velasco-Berrelleza, V., Piperakis, M. M., Mitchenall, L. A., Cugliandolo, F. M., Beton, J. G., Stevenson, C. E. M., Hoogenboom, B. W., Bates, A. D., Maxwell, A., and Harris, S. A. (2021) Base-pair resolution analysis of the effect of supercoiling on DNA flexibility and major groove recognition by triplex-forming oligonucleotides. Nat. Commun., 12, 1053.

[25] Lankaš, F., Lavery, R., and Maddocks, J. H. (2006) Kinking Occurs during Molecular Dynamics Simulations of Small DNA Minicircles. Structure, 14, 1527–1534.

[26] Mitchell, J. S., Laughton, C. A., and Harris, S. A. (2011) Atomistic simulations reveal bubbles, kinks and wrinkles in supercoiled DNA. Nucleic Acids Res., 39, 3928–3938.

[27] Matek, C., Ouldridge, T. E., Doye, J. P. K., and Louis, A. A. (2015) Plectoneme tip bubbles: Coupled denaturation and writhing in supercoiled DNA. Sci. Rep., 5, 7655.

[28] Amzallag, A., Vaillant, C., Jacob, M., Unser, M., Bednar, J., Kahn, J. D., Dubochet, J., Stasiak, A., and Maddocks, J. H. (2006) 3D reconstruction and comparison of shapes of DNA minicircles observed by cryo-electron microscopy. Nucleic Acids Research, 34, e125.

[29] Irobalieva, R. N., Fogg, J. M., Catanese, D. J., Sutthibutpong, T., Chen, M., Barker, A. K., Ludtke, S. J., Harris, S. A., Schmid, M. F., Chiu, W., and Zechiedrich, L. (2015) Structural diversity of supercoiled DNA. Nat. Commun., 6, 1–10.

[30] Wei, J., Czapla, L., Grosner, M. A., Swigon, D., and Olson, W. K. (2014) DNA topology confers sequence specificity to nonspecific architectural proteins. Proc. Natl. Acad. Sci. U.S.A., 111, 16742–16747.

[31] Pasi, M. and Lavery, R. (2016) Structure and dynamics of DNA loops on nucleosomes studied with atomistic, microsecond-scale molecular dynamics. Nucleic Acids Res., 44, 5450–5456.

[32] Pasi, M., Mornico, D., Volant, S., Juchet, A., Batisse, J., Bouchier, C., Parissi, V., Ruff, M., Lavery, R., and Lavigne, M. (2016) DNA minicircles clarify the specific role of DNA structure on retroviral integration. Nucleic Acids Res., 44, 7830–7847.

[33] D’Annessa, I., Coletta, A., Sutthibutpong, T., Mitchell, J., Chillemi, G., Harris, S., and Desideri, A. (2014) Simulations of DNA topoisomerase 1B bound to supercoiled DNA reveal changes in the flexibility pattern of the enzyme and a secondary protein-DNA binding site. Nucleic Acids Res., 42, 9304–9312.

[34] Noy, A., Maxwell, A., and Harris, S. A. (2017) Interference between Triplex and Protein Binding to Distal Sites on Supercoiled DNA. Biophys. J., 112, 523–531.

[35] Fogg, J. M., Randall, G. L., Pettitt, B. M., Sumners, D. W. L., Harris, S. A., and Zechiedrich, L. (2012) Bullied no more: when and how DNA shoves proteins around. Q. Rev. Biophys., 45, 257–299.

[36] Teter, B., Goodman, S. D., and Galas, D. J. (2000) DNA bending and twisting properties of integration host factor determined by DNA cyclization. Plasmid, 43, 73–84.

[37] Bao, Q., Chen, H., Liu, Y., Yan, J., Dröge, P., and Davey, C. A. (2007) A Divalent Metal-mediated Switch Controlling Protein-induced DNA Bending. J. Mol. Biol., 367, 731–740.

[38] Richet, E., Abcarian, P., and Nash, H. A. (1986) The interaction of recombination proteins with supercoiled DNA: Defining the role of supercoiling in lambda integrative recombination. Cell, 46, 1011–1021.

[39] Normanno, D., Vanzi, F., and Pavone, F. S. (2008) Single-molecule manipulation reveals supercoiling-dependent modulation of lac repressor-mediated DNA looping. Nucleic Acids Res., 36, 2505–2513.

[40] Dorman, C. J. and Ní Bhriain, N. (2020) CRISPR-Cas, DNA Supercoiling, and Nucleoid-Associated Proteins. Trends in Microbiol., 28, 19–27.

[41] Reverchon, S., Meyer, S., Forquet, R., Hommais, F., Muskhelishvili, G., and Nasser, W. (12, 2020) The nucleoid-associated protein IHF acts as a ‘transcriptional domainin’ protein coordinating the bacterial virulence traits with global transcription. Nucleic Acids Res., 49, 776–790.

[42] Case, D. A. et al. AMBER. (2016) v16.

[43] Fogg, J. M., Kolmakova, N., Rees, I., Magonov, S., Hansma, H., Perona, J. J., and Zechiedrich, E. L. (2006) Exploring writhe in supercoiled minicircle DNA. J. Phys. Cond. Matt., 18, S45.

[44] Nguyen, H., Roe, D. R., and Simmerling, C. (2013) Improved generalized born solvent model parameters for protein simulations. J. Chem. Theor. Comput., 9, 2020–2034.

[45] Perez, A., MacCallum, J. L., Brini, E., Simmerling, C., and Dill, K. A. (2015) Grid-Based Backbone Correction to the ff12SB Protein Force Field for Implicit-Solvent Simulations. J. Chem. Theor. Comput., 11, 4770–4779.

[46] Nguyen, H., Pérez, A., Bermeo, S., and Simmerling, C. (2015) Refinement of Generalized Born Implicit Solvation Parameters for Nucleic Acids and Their Complexes with Proteins. J. Chem. Theor. Comput., 11, 3714–3728.

[47] Anandakrishnan, R., Drozdetski, A., Walker, R. C., and Onufriev, A. V. (2015) Speed of conformational change: comparing explicit and implicit solvent molecular dynamics simulations.. Biophys. J, 108, 1153–1164.

[48] Maier, J. A., Martinez, C., Kasavajhala, K., Wickstrom, L., Hauser, K. E., and Simmerling, C. (2015) ff14SB: Improving the Accuracy of Protein Side Chain and Backbone Parameters from ff99SB. J. Chem. Theor. Comput., 11, 3696–3713.

[49] Ivani, I., Dans, P. D., Noy, A., Pérez, A., Faustino, I., Hospital, A., Walther, J., Andrio, P., Goñi, R., Balaceanu, A., Portella, G., Battistini, F., Gelpí, J. L., González, C., Vendruscolo, M., Laughton, C. A., Harris, S. A., Case, D. A., and Orozco, M. (2015) Parmbsc1: A refined force field for DNA simulations. Nat. Meth., 13, 55–58.

[50] Sutthibutpong, T., Harris, S. A., and Noy, A. (2015) Comparison of Molecular Contours for Measuring Writhe in Atomistic Supercoiled DNA. J. Chem. Theor. Comput., 11, 2768–2775.

[51] Britton, L. A., Olson, W. K., and Tobias, I. (2009) Two perspectives on the twist of DNA. J. Chem. Phys., 131, 245101.

[52] Lu, X. and Olson, W. K. (2003) 3DNA: a software package for the analysis, rebuilding and visualization of three-dimensional nucleic acid structures. Nucleic Acids Res., 31, 5108–5121.

[53] Velasco-Berrelleza, V., Burman, M., Shepherd, J. W., Leake, M. C., Golestanian, R., and Noy, A. (2020) SerraNA: a program to determine nucleic acids elasticity from simulation data. Phys. Chem. Chem. Phys., 22, 19254–19266.

[54] Dans, P. D., Faustino, I., Battistini, F., Zakrzewska, K., Lavery, R., and Orozco, M. (2014) Unraveling the sequence-dependent polymorphic behavior of d(CpG) steps in B-DNA. Nuc. Acids Res., 42, 11304–11320.

[55] Roe, D. R. and Cheatham, T. E. (2013) PTRAJ and CPPTRAJ: Software for Processing and Analysis of Molecular Dynamics Trajectory Data. J. Chem. Theor. Comput., 9, 3084–3095.

[56] Sugimura, S. and Crothers, D. M. (2006) Stepwise binding and bending of DNA by Escherichia coli integration host factor. Proc. Natl. Acad. Sci. USA, 103, 18510–18514.

[57] Wang, Q., Irobalieva, R. N., Chiu, W., Schmid, M. F., Fogg, J. M., Zechiedrich, L., and Pettitt, B. M. (2017) Influence of DNA sequence on the structure of minicircles under torsional stress. Nucleic Acids Res., 45, 7633–7642.

[58] Sutthibutpong, T., Matek, C., Benham, C., Slade, G. G., Noy, A., Laughton, C., Doye, J. P., Louis, A. A., and Harris, S. A. (2016) Long-range correlations in the mechanics of small DNA circles under topological stress revealed by multi-scale simulation. Nucleic Acids Res., 44, 9121–9130.

[59] Lal, A., Dhar, A., Trostel, A., Kouzine, F., Seshasayee, A. S. N., and Adhya, S. (2016) Genome scale patterns of supercoiling in a bacterial chromosome. Nat. Commun., 7, 11055.

[60] Hardy, C. D. and Cozzarelli, N. R. (2005) A genetic selection for supercoiling mutants of Escherichiacoli reveals proteins implicated in chromosome structure. Molecular Microbiol., 57, 1636–1652.

[61] Desai, P. R., Brahmachari, S., Marko, J. F., Das, S., and Neuman, K. C. (2020) Coarse-grained modelling of DNA plectoneme pinning in the presence of base-pair mismatches. Nuc. Acids Res., 48, 10713–10725.

[62] Lim, W., Randisi, F., Doye, J. P. K., and Louis, A. A. (2022) The interplay of supercoiling and thymine dimers in DNA. Nucleic Acids Res.,.

[63] Norregaard, K., Andersson, M., Sneppen, K., Nielsen, P. E., Brown, S., and Oddershede, L. B. (2013) DNA supercoiling enhances cooperativity and efficiency of an epigenetic switch. Proc. Natl. Acad. Sci. U.S.A., 110, 17386–17391.

[64] Yan, Y., Xu, W., Kumar, S., Zhang, A., Leng, F., Dunlap, D., and Finzi, L. (11, 2021) Negative DNA supercoiling makes protein-mediated looping deterministic and ergodic within the bacterial doubling time. Nucleic Acids Res., 49, 11550–11559.

[65] Fisher, G. L., Pastrana, C. L., Higman, V. A., Koh, A., Taylor, J. A., Butterer, A., Craggs, T., Sobott, F., Murray, H., Crump, M. P., Moreno-Herrero, F., and Dillingham, M. S. (2017) The structural basis for dynamic DNA binding and bridging interactions which condense the bacterial centromere. eLife, 6, e28086.

[66] Ding, Y., Manzo, C., Fulcrand, G., Leng, F., Dunlap, D., and Finzi, L. (2014) DNA supercoiling: A regulatory signal for the lambda repressor. Proc. Natl. Acad. Sci., 111, 15402–15407.

[67] Yan, Y., Ding, Y., Leng, F., Dunlap, D., and Finzi, L. (2018) Protein-mediated loops in supercoiled DNA create large topological domains. Nucleic Acids Res., 46, 4417–4424.

